# Combination Treatment with NCX 4040 and Napabucasin Triggers Enhanced Oxidative Stress, STAT3 Inhibition and Plausible Immunogenic Activation in Metastatic Prostate Cancer Cells

**DOI:** 10.1101/2025.09.09.675191

**Authors:** Nishtha Pathak, Shrushti Shah, Jeffrey Mathew, Gnanasekar Munirathinam

## Abstract

Prostate cancer (PCa) remains a significant health concern, ranking as the second most common cancer after lung cancer, despite recent advances in diagnostics and therapeutics. Due to its heterogeneous nature, diverse etiology, and the limited efficacy of current treatments, developing a therapeutic approach that can address the multifaceted aspects of the disease is imperative. Combinatorial therapy has emerged as a promising strategy for addressing tumor heterogeneity and the limitations of current treatments across various cancer types.

In our study, we investigated the potential of combining NCX 4040, a nitric oxide-releasing aspirin derivative, and Napabucasin, a STAT3 and cancer stemness inhibitor, as a novel PCa treatment strategy. We utilized two cellular models: BPH-Cd, a cadmium-transformed carcinogenic cell line, and DU145, a brain metastatic PCa cell line. Our findings demonstrate that the combination treatment exerted a synergistic and dose-dependent reduction in cell viability, tumorigenicity, and migratory capabilities in both BPH-Cd and DU145 cells, surpassing the effects observed with individual treatments. Furthermore, this combination treatment triggered a robust generation of cellular and mitochondrial reactive oxygen species (ROS), resulting in G2/M cell cycle arrest and late-stage apoptosis in both cell lines.

Additionally, the combination treatment downregulated redox-sensitive transcription factors pSTAT3 Tyr705 and Ser727, as well as pro-survival proteins TRX1/2, TRXR1, and GPX4, while upregulating tumor suppressor proteins PRDX1 and TXNIP. This implies that inhibition of the pro-survival redox proteins leads to the accumulation of oxidized PRDX1 and TXNIP, thereby elevating oxidative stress and blocking STAT3-mediated transcription, ultimately impeding cell proliferation and survival. Moreover, increased expression of pH2AX protein, a DNA damage marker, indicated that the combination induced DNA damage, resulting in the activation of the cGAS-STING pathway, an anti-tumor immunogenic mechanism as confirmed with immunoblotting. Downregulation of various stem cell markers linked to Wnt/β-catenin and Notch-1 signaling pathways highlight the combination’s ability to also target cancer stemness. In summary, our study underscores the promise of combining NCX 4040 and Napabucasin as an innovative and multifaceted therapeutic approach for PCa.

**Abstract Figure:** 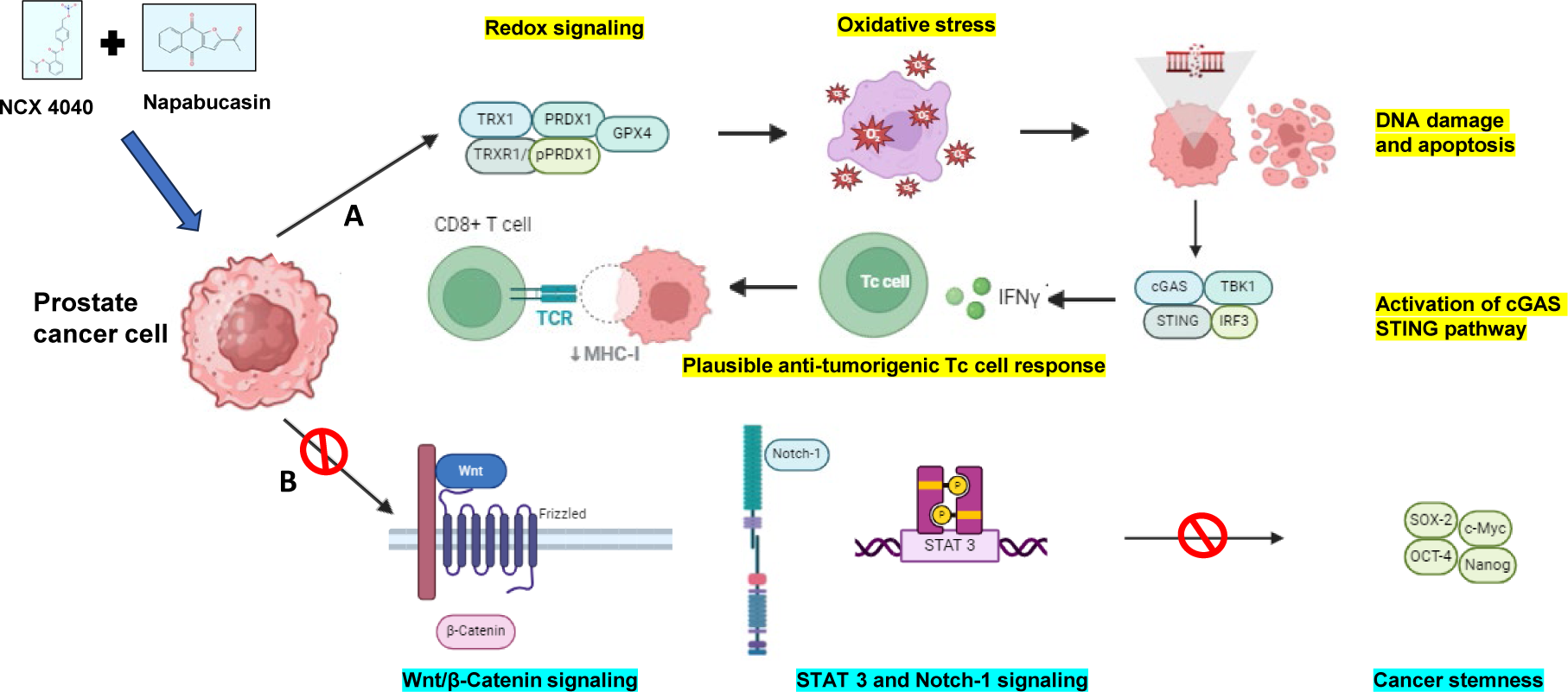

(A) The combination therapy modulates key markers of the redox signaling pathway, inducing excessive oxidative stress in PCa cells, which subsequently leads to DNA damage and late apoptotic cell death. DNA damage activates the cGAS-STING pathway, upregulating interferon regulatory factor 3 (IRF3), which may, in turn, trigger the activation of tumor-infiltrating lymphocytes, generating a potent anti-tumorigenic response. (B) The combination therapy inhibits stem-cell signaling pathways, including Wnt/β-catenin, Notch-1, and STAT3, potentially suppressing cancer stemness and contributing to long-term disease control.

## Introduction

Prostate Cancer (PCa) stands as a prevalent non-cutaneous malignancy affecting the male population in the United States and ranks as the second leading cause of cancer-related morbidity, closely following lung cancer.^1^ PCa exhibits pronounced morphological heterogeneity and is notorious for its multifocality, varying capacity for metastatic seeding, and inherent treatment resistance. While various treatment modalities, including radiation, androgen deprivation therapy (ADT),^2^ chemotherapy,^3^ and immunotherapy,^4^ are currently available for PCa management, their efficacy is limited, particularly in high-risk and metastatic subtypes. Patients with advanced PCa often develop castration resistance or experience relapses having only 15% survival rate, highlighting the need for comprehensive therapeutic strategies.^5^

Given the inherent heterogeneity of PCa, which encompasses a diverse molecular landscape that remains under investigation, addressing this complex disease requires a multifaceted therapeutic approach. Monotherapy with a single drug has proven insufficient in tackling this nature of PCa, necessitating exploration of combination therapies involving two or more agents. Combinatorial therapy has emerged as a promising avenue for various cancer types, offering the potential to target diverse facets of the disease and overcome treatment limitations.^6–8^ Hence, in this study, we investigated the synergistic and combinatorial effects of two drug compounds, NCX 4040 and Napabucasin, in the context of PCa therapy.

NCX 4040, a nitric oxide-releasing non-steroidal anti-inflammatory drug (NO-NSAID), is a derivative of aspirin, renowned for its potent cytolytic effects against various cancer types.^9^ Existing evidence supports a plausible mechanism of action wherein the drug induces oxidative stress by depleting cellular glutathione (GSH), resulting in the generation of reactive oxygen species (ROS) or quinone methide, culminating in DNA damage and cell death.^10^ Our prior investigations have also established that NCX 4040 provokes oxidative stress via hydrogen peroxide, leading to mitochondrial-mediated cell death in the metastatic PC3 cell line.^11^ Consequently, the current body of research underscores the remarkable cytotoxic potential of NCX 4040.^12–14^ Furthermore, the structural spacer moiety in NCX 4040 presents an intriguing avenue for exploration, with the possibility of assessing its synergistic potential when combined with Napabucasin. This endeavor holds promise for developing a highly potent therapeutic strategy against advanced PCa.

On the other hand, Napabucasin (BBI608) stands out as a recently developed small molecule inhibitor targeting the signal transducer and activator of transcription 3 (STAT3) transcription factor.^15^ It has exhibited its potential in inhibiting self-renewal and triggering apoptosis in cancer stem cells across a range of cancer types, including glioblastoma,^16^ colorectal,^17^ pancreatic,^18^ non-small cell lung,^19^ and gastric cancer.^20^ Notably, this orally administered agent has displayed remarkable anti-tumor and anti-metastatic properties both as a standalone treatment and in synergy with various chemotherapeutic drugs, such as paclitaxel, as confirmed by in-vivo studies.^7, 21–24^

Moreover, research has shed light on Napabucasin’s role in fostering immunogenic cell death, with demonstrated effects on the expression of calreticulin—a prominent “eat me” molecule. This, in turn, leads to STAT3 inhibition and the activation of dendritic cells, facilitating increased infiltration of CD4+ and CD8+ T cells.^25^ Additionally, it has been proposed that Napabucasin induces a substantial rise in reactive oxygen species (ROS) and provokes DNA damage, resulting in a STAT3 phosphorylation decrease that is dependent on NQO1 and ROS.^26^ Consequently, Napabucasin presents a promising candidate for combination therapy, effectively addressing multifaceted aspects of cancer. Notably, the potential of Napabucasin in the context of PCa remains largely unexplored. Limited studies demonstrate its high efficacy and anti-cancer activity in bone metastasis,^27^ cancer stem cell microtumors^28^ and castration-resistant PCa (CRPC).^29^ Thus, investigating its efficacy in suppressing PCa cell proliferation, migration, and stemness holds promise for opening new avenues in cancer patient care.

Building upon this, our research aims to shed light on the potential of this combination therapy approach to address the unique challenges posed by PCa, with the ultimate goal of advancing the development of potent therapeutics for this malignancy.

## Materials and Methods

### Cell Culture

BPH-Cd cell line was developed by our lab as a cellular model for recapitulating cadmium induced prostate carcinogenesis. DU145 was bought from ATCC and BPH-1 was obtained from Dr. Hayward, NorthShore University HealthSystem. All the cell lines were cultured in complete Roswell Park Memorial Institute (RPMI) 1640 (Hyclone) medium, supplemented with 10% fetal bovine serum (FBS) (Corning), 1% 100X antimycotic (Sigma Aldrich) and 100 μL 30 mg/mL gentamycin (MP Biomedicals) in a 5% CO2 incubator at 37°C. BPH-Cd cells were additionally supplemented with 10 μM cadmium chloride (Sigma Aldrich) solution to maintain the cadmium carcinogenicity.

### Chemicals

NCX 4040 (331.28 g/mol) was purchased from Tocris Bioscience. Napabucasin (2-Acetylnaphtho[2,3-b] furan-4,9-dione - 240.21 g/mol) was purchased from TCI Organic Chemicals. pSTAT3 (Tyr705), pSTAT3 (Ser727), redox homeostasis and signaling antibody sampler kit, STING pathway antibody sampler kit, pH2AX, stem cell marker antibodies, β-actin, anti-rabbit IgG and anti-mouse IgG secondary antibody were purchased from Cell Signaling Technology. Thiazolyl Blue Tetrazolium Bromide (MTT Dye), crystal violet dye, Dichlorodihydrofluorescein Diacetate (DCFH-DA), 4ʹ,6-diamidino-2-phenylindole (DAPI) were obtained from Sigma Aldrich. FITC Annexin V Apoptosis Detection Kit and Propidium Iodide were obtained from BD Biosciences. Calcein AM green dye and MitoSOX red were purchased from Invitrogen.

### Cell Viability Assay

BPH-Cd and DU145 cells were seeded at a seeding density of 3000 and 4500 cells/well in 96-well plates respectively and incubated at 37°C in a 5% CO2 incubator, until 50% confluency was attained. The cells were then treated with different concentrations of NCX 4040 and Napabucasin alone and in combination for 48 hours. After 48 hours of drug treatment, 0.5 mg/mL MTT dye was added to each well and the plates were then incubated for 2-3 hours. The crystals were then dissolved by adding the solubilizing agent and the absorbance was then measured at 570 nm, followed by calculating the cell viability.^30^

### Clonogenic Assay

500-700 cells were seeded in each well of 6-well plate. The plates were incubated at 37°C in a 5% CO_2_ incubator until the cells had formed visible number of colonies. The cells were then treated with various treatment groups for 5-7 days. Following treatment, the cells were fixed using a fixative solution containing 1 part of acetic acid and 7 parts of methanol. The fixed cells were stained with 0.5% crystal violet dye and for 2 hours and the number of colonies were manually counted.^11^

### Spheroid Assay

BPH-Cd cells were seeded with a density of 1500 cells/well in Matrigel coated 96 well plate. After significant growth of spheroids, they were treated with various concentrations of the drugs alone and in combination for 4-5 days. The treated spheroids were then stained with 10 μM Calcein-AM green dye for 30 minutes at 37°C in a 5% CO_2_ incubator and the images were recorded by using Keyence microscope.^31^

### Wound Healing Assay

BPH-Cd and DU145 cells were seeded at a seeding density of 2 x 10^5^ cells/well in 24 well plates. Upon 100% confluency, a scratch was made at the center of each well and the cells were treated. The images were recorded at 0, 24 and 48 hours. Gap closure was evaluated, and the percentage of wound healing ability was measured by analyzing the wound closure using T-Scratch software.^11^

### Transwell Assay

Cell migration was performed by using 24 well Transwell chambers (Millipore, Billerica, MA, USA). Approximately 1.5 x 10^4^ BPH-Cd and DU145 cells were seeded in each insert resuspended in incomplete media and subsequently treated with varying drug concentrations for 48 hours. Complete RPMI with 10% FBS was used as a chemoattractant in the bottom chamber. After treatment, the migrated cells at the bottom of the insert were stained with Hema 3 Stat Pack staining kit and the migrated cells were imaged using a bright field microscope. ^11^

### Cellular ROS Analysis

BPH-Cd and DU145 cells were seeded at a seeding density of 1.5 x 10^5^ cells in 6-well plates and incubated at 37°C in a 5% CO_2_ incubator. Upon 60-70% confluency, the cells were stained with 20 μM of DCFH-DA dye, prepared in phenol red-free RPMI with 1% FBS, for 45 minutes. The samples were then washed and treated for 15-20 minutes with varying concentrations of the drugs prepared in the same media. Hydrogen peroxide served as a positive control. Cellular ROS was analyzed using a flow cytometer.^11^

### Mitochondrial ROS Analysis

1.5 x 10^5^ cells were seeded in 6-well plates and upon 60-70% confluency, the cells were treated with varying concentrations of the drugs prepared in phenol red free RPMI with 1% FBS for 1.5 hours. Later, the cells were then stained with 5 μM of mitoSOX red stain for 30 min at RT. Mitochondrial ROS was analyzed using a flow cytometer.^32^

### Cell Cycle Analysis

BPH-Cd and DU145 cells were seeded at a seeding density of 1.5 x 10^5^ cells in a T25 flasks and incubated at 37°C in a 5% CO_2_ incubator. Once the cells were approximately 50% confluent, they were treated with varying concentrations of NCX 4040 and Napabucasin for 48 hours. Later, the cells were fixed overnight with ice-cold 70% ethanol at 4°C. The next day, the cells were treated with RNAase I and then stained with propidium iodide (PI), prepared as per manufacturer’s instruction, for 30 minutes at RT. The samples were analyzed using a flow cytometer.^33^

### Cell Death Analysis

1.5 x 10^5^ cells were seeded in 6-well plates and upon 50% confluency, the cells were serum starved overnight. Later, the cells were treated with varying concentrations of NCX 4040 and Napabucasin for 48 hours. Both live and dead cells were pooled together and were stained with Annexin V-FITC Apoptosis Detection Kit as per the manufacturer’s instruction. The samples were analyzed using a flow cytometer.^11^

### Immunofluorescence Analysis

BPH-Cd and DU145 cells were seeded at a seeding density of 6 x 10^3^ cells/well in an 8-well chamber slide and incubated at 37°C in 5% CO_2_ incubator. Once the cells were 70% confluent, they were treated with varying concentrations of NCX 4040 and Napabucasin alone and in combination for 48 hours. Following the treatment, the cells were fixed using 4% paraformaldehyde for 20 minutes, permeabilized with 0.1% triton X-100 for 10 minutes, followed by blocking them with 5% normal goat serum blocking buffer for 1 hour at RT. The cells were incubated with primary antibody (at 1:250 dilution) specific to phospho-H2AX (γH2AX or pH2AX) overnight at 4°C, followed by 1 hour incubation at RT with FITC labelled secondary antibody. The cells were then stained with DAPI and the slide was mounted using a fluorogel mounting medium. The images were taken at 60X in the confocal fluorescent microscope.^34^

### Western Blot Analysis

T-25 flasks were seeded with cells and incubated at 37°C with 5% CO_2_ until 70% confluent. Later the cells were treated with varying drug concentrations for 48 hours. For detection of phosphorylated proteins, other flasks were serum starved overnight by replacing the complete media with serum-free incomplete media upon 50-60% confluency, followed by the same treatments the next day. The cell lysate was prepared by using Mammalian Protein Extraction Reagent (M-PER – ThermoFisher Scientific) as per the manufacturer’s instructions. Protein concentration was evaluated using rapid gold BCA assay and SDS-PAGE was performed followed by transfer to nitrocellulose membrane. The blots were blocked with 5% skimmed milk and probed with respective primary antibodies purchased from Cell Signaling Technology. Later, the blots were probed with corresponding secondary antibodies and specific protein expression was detected by using SuperSignal West Pico Plus Chemiluminescent Substrate (ThermoFisher Scientific) as per the manufacturer’s instruction. The blots were visualized on a ChemiDoc system and developed on X-Ray films. Band intensities were then analyzed using ImageJ Software.^30^

### Statistical Analysis

Statistical analysis was performed using multi-variate Tuckey and Dunnett tests to determine significance of every individual treatment group to their respective combinations and every treatment group with the untreated control sample respectively. P values less than 0.05 were considered significant. The sample size for all the experiments was n=3. SAS 9.4 was used to perform the statistical analysis for each experiment.

## Results

### NCX 4040 synergizes with the effects of Napabucasin in inhibiting cell viability

We first evaluated the synergy between NCX 4040 and Napabucasin to determine their combinatorial anti-cancer effect in BPH-Cd and DU145 cells. The cells were exposed to varied concentrations of NCX 4040 (2.5 μM, 5 μM, and 10 μM) and Napabucasin (100 nM and 250 nM), either alone or in combination. The synergistic effects were assessed by comparing the cytotoxic effects of individual and combined treatments and calculating the synergy score. Both BPH-Cd and DU145 cells displayed a decline in percent cell viability in a dose-dependent manner as shown in Figure 1A and 1B. Analysis of synergy scores unveiled a synergistic interaction between NCX 4040 and Napabucasin (as depicted in Figure 1C, 1D and 1E), which explained the significant reduction in percent cell viability in the combined treatment, in contrast to individual treatments. Notably, two outliers were identified: the combination of NCX 2.5 μM and NAPA 250 nM exhibited an antagonistic effect, while NCX 5 μM plus NAPA 250 nM demonstrated an additive effect in BPH-Cd cells (see Figure 1C). Subsequently, the former combination was excluded from further exploration in subsequent experiments. Furthermore, when treating BPH-1 cells, a non-carcinogenic cell line, our results indicated that the combination treatment was relatively less toxic to normal cells, as there was no significant reduction in percent cell viability (shown in Figure 1F). This suggests a potential selectivity of the combination treatment towards cancer cells while sparing normal cells.

**Figure 1:**
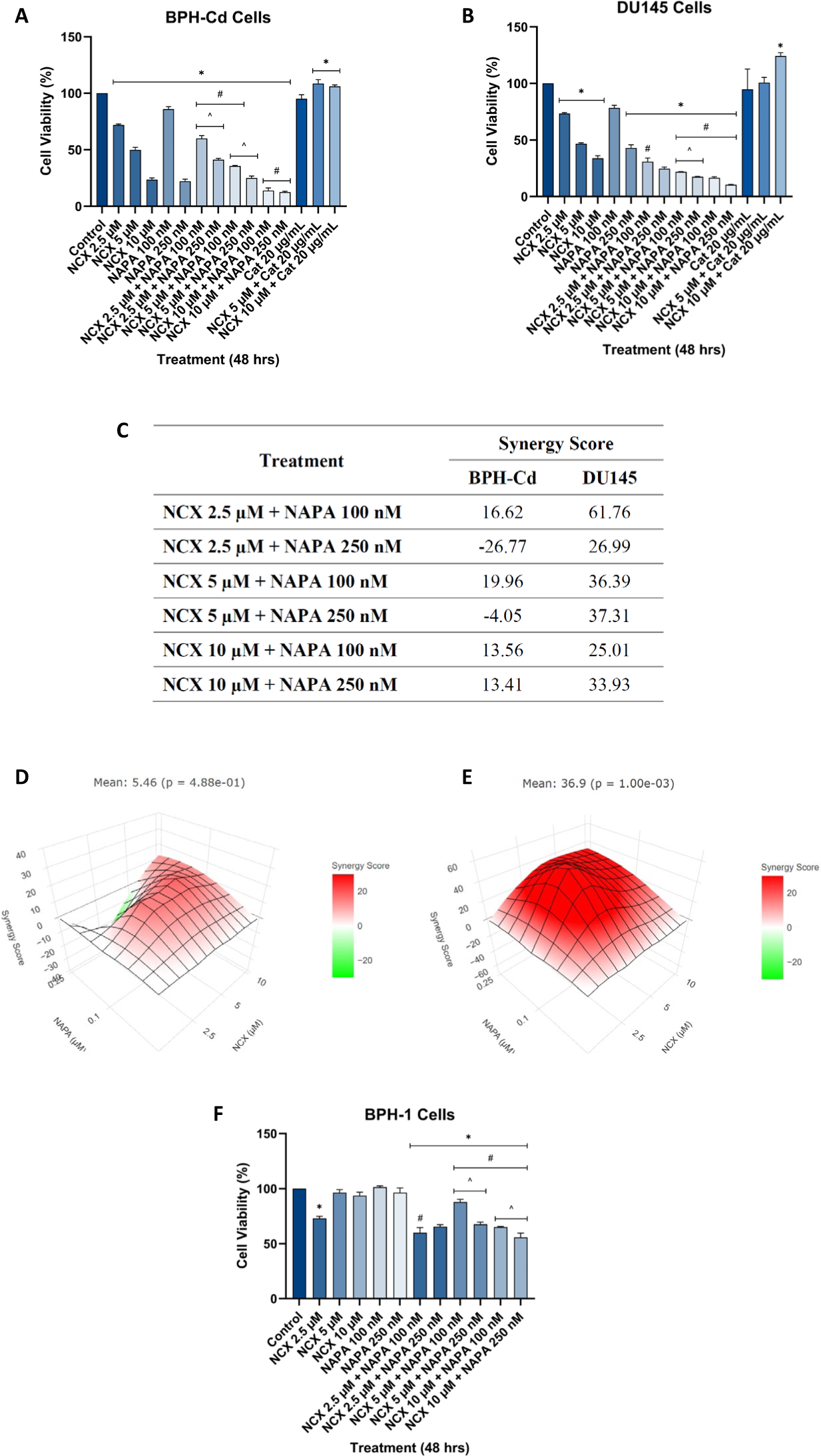
NCX 4040 synergizes with the effects of Napabucasin in inhibiting cell viability of A) BPH-Cd and B) DU145 cells. MTT assay was performed on cells treated with different concentrations of NCX 4040 and Napabucasin, alone and in combination, for 48 hours. Catalase, an antioxidant, was used as an inhibitor to assess if it can block the anti-cancer activity of NCX 4040. The combination treatment effectively inhibited cell viability in a dose-dependent manner, as compared to the alone treatments. The IC50 values were found to be 5 μM for NCX 4040 and 250 nM for Napabucasin. Catalase blocked the anti-cancer activity of NCX 4040, which implied that oxidative stress could be the mechanism of action. **C)** Synergy scores of NCX 4040 and Napabucasin drug combination calculated using Highest Single Agent (HSA) model in synergyFinder+ web tool. NCX 4040 and Napabucasin are shown to exhibit moderate to high synergistic effects in **D)** BPH-Cd and **E)** DU145 cells. **F)** Safety profile base of off MTT assay on BPH-1 cells demonstrated relatively low levels of cytotoxicity, ensuring minimal off-target effects. Data are means ± SD (n=3). * p < 0.05: compared to control; # p < 0.05: mean of combination treatment compared to their respective individual treatments; and ^ p < 0.05: mean of combination treatment compared to other combination treatments.

### NCX 4040 and Napabucasin impeded the in-vitro tumorigenic potential and migratory properties of PCa cells

Next, we evaluated the potential of combination treatment in inhibiting the tumorigenic potential and migratory properties of PCa cells. The combined treatment exhibited a heightened suppression of in-vitro tumorigenic potential in both cell lines, as evidenced by clonogenic and spheroid assays in Figure 2. In contrast to the outcomes observed in the individual treatment groups, the concurrent application of NCX 4040 and Napabucasin induces a substantial reduction in colony numbers for both BPH-Cd and DU145 cells (see Figures 2A-2D), along with a notable decrease in the size and quantity of spheroids in BPH-Cd cells (see Figures 2E and 2F). These findings suggest that the combined treatment proves more efficacious in impeding tumorigenic potential and in-vitro tumor formation within a 3D microenvironment.

**Figure 2:**
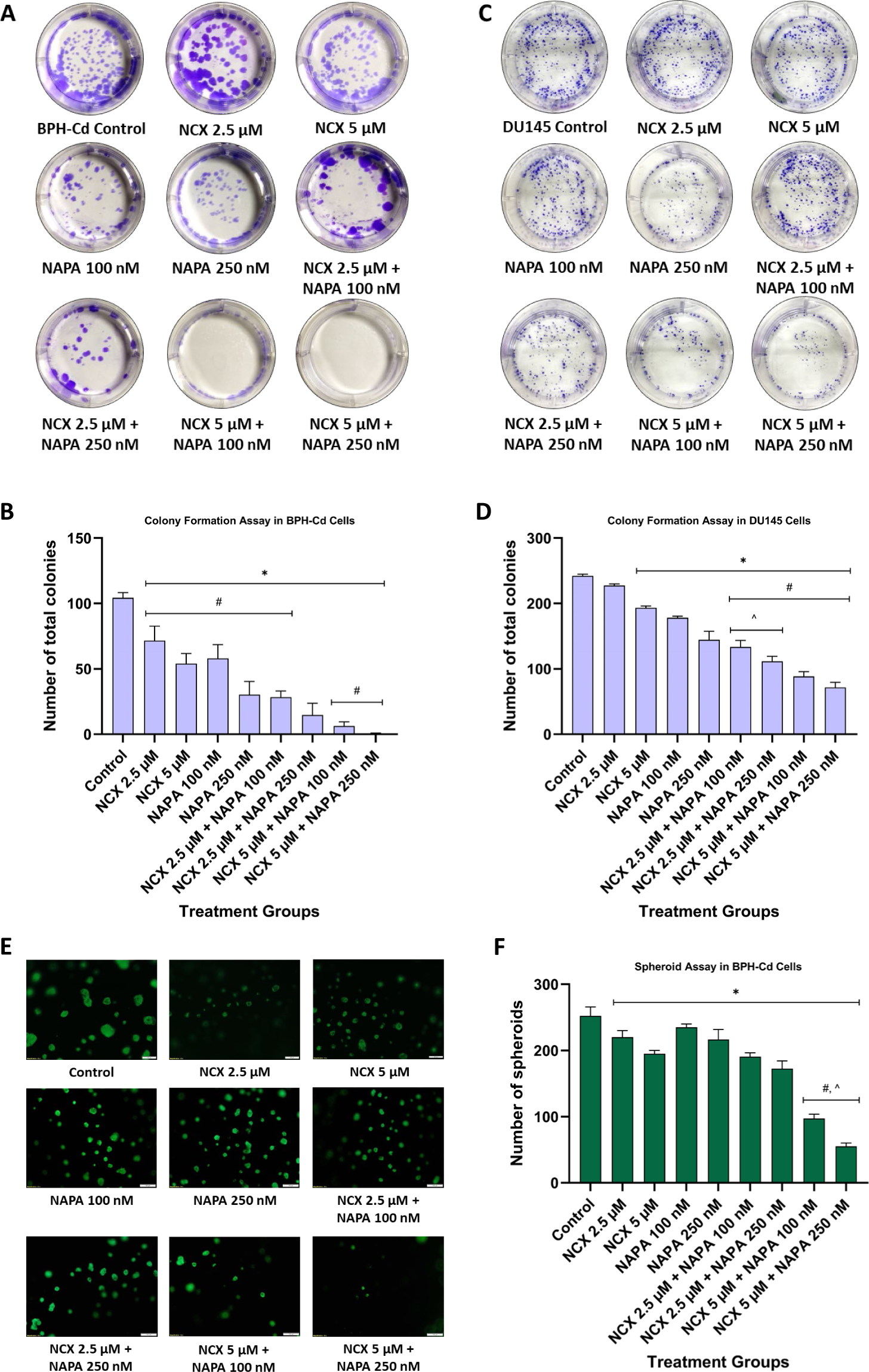
NCX4040 in combination with Napabucasin synergistically inhibits the tumorigenic potential of PCa cells. To evaluate the anti-tumorigenic potential of the combination treatment, a clonogenic assay was performed in **A)** BPH-Cd and **C)** DU145 cells. Quantitative analysis of clonogenic assay is represented in figures **B)** and **D)** respectively. The colonies of cells were treated with varying concentrations of both drugs, alone and in combination, for 4-5 days. The experiment was terminated by staining the colonies with crystal violet and quantifying the number of colonies manually. A dose-dependent reduction in the size and number of colonies was observed, with the combination treatment proving to be more effective than individual treatments in inhibiting the clonogenic potential of both cell lines. **E)** In 3D microenvironment, the spheroid assay depicted dose-dependent reduction in the size and number of spheroids was observed, with the combination treatment proving to be more effective than individual treatments in inhibiting the tumorsphere formation potential of BPH-Cd cells. **F)** represents the quantitative analysis of spheroid assay in BPH-Cd cells. These findings suggest that the combination treatment has potent inhibitory effects on in-vitro tumorigenic potential of the cancer cells in both 2D and 3D environments. Data are means ± SD (n=3). * p < 0.05: compared to control; # p < 0.05: mean of combination treatment compared to their respective individual treatments; and ^ p < 0.05: mean of combination treatment compared to other combination treatments.

Moreover, the combined treatment markedly suppresses the migratory capabilities of both cell lines, resulting in approximately 90% and 75% inhibition in BPH-Cd and DU145 cells, respectively, after 48 hours compared to their respective controls. Time-dependent wound healing assays conducted at 0-, 24-, and 48-hours post-treatment demonstrate a reduced open wound area and accelerated migration in the individual treatment groups, as shown in Figure 3.

**Figure 3:**
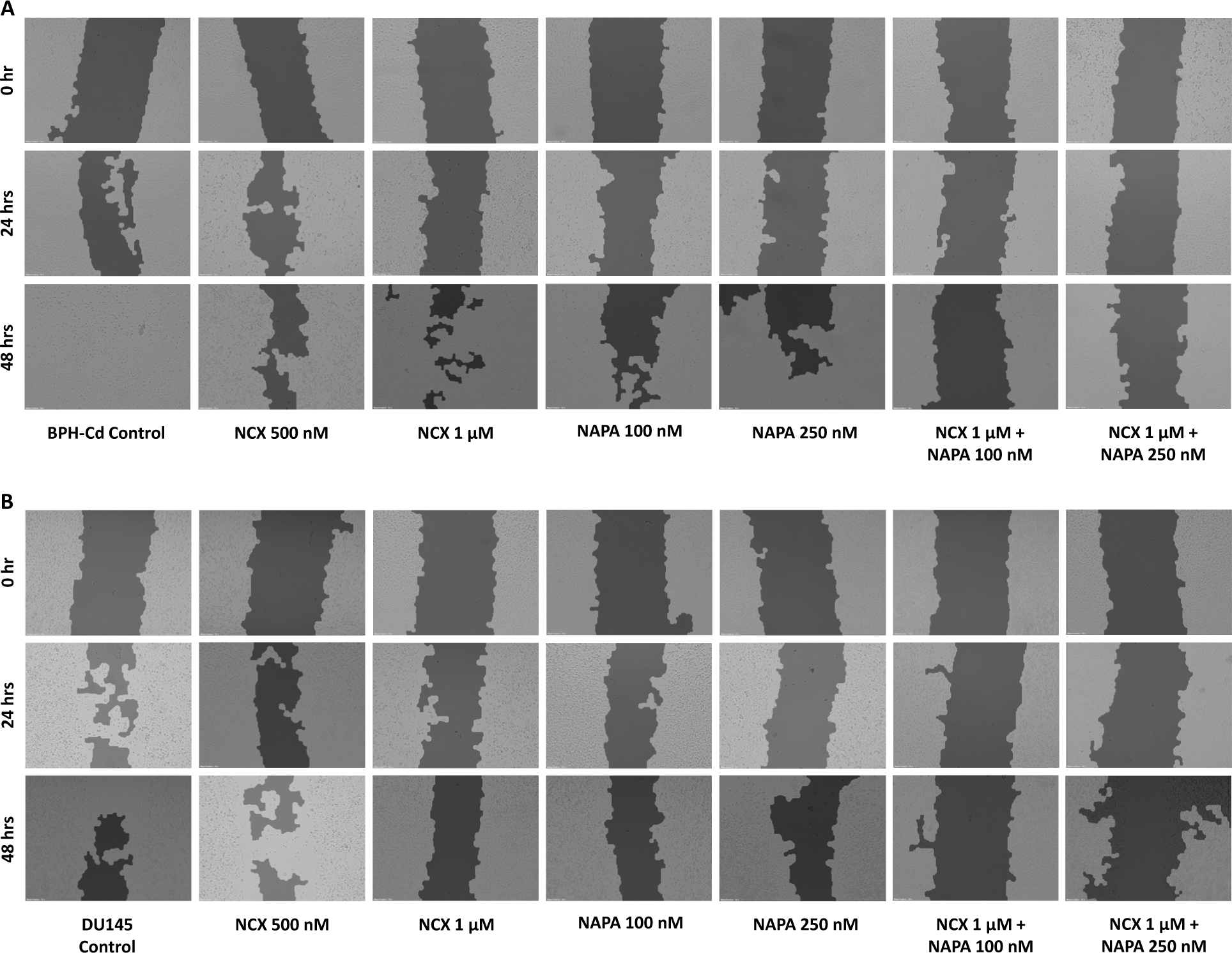
Combination treatment inhibits the migration potential of A) BPH-Cd and B) DU145 cells. The wound healing assay was carried out to investigate the effect of combination treatment on the wound healing ability of cancer cells. A scratch was then made in each well when the cells reached 100% confluency and the cells were exposed to varying concentrations of both drugs, either alone or in combination. Images were captured at 0, 24, and 48 hours to monitor the migration of cells and the wound closure process. As shown, the combination treatment significantly inhibited the migration of cells towards the open area, resulting in a complete halt of wound closure in a dose and time dependent manner. This indicates that the combination treatment is highly effective in suppressing the metastatic properties of these cells, as evidenced by the reduced wound healing ability. Data is representative of one of three experiments (N=3). Scale bar 100 μm. Magnification 10X.

### NCX 4040 in combination with Napabucasin induces cellular and mitochondrial ROS production leading to G2/M cell cycle arrest and late apoptosis

Building upon prior research from our laboratory, we were aware that NCX 4040 induces ROS production, leading to oxidative stress-related cellular death in PC3 cells.^11^ Conversely, existing literature indicates that Napabucasin acts as a substrate for intracellular oxidoreductases, such as NQO1, which are highly expressed in cancer cells. This interaction ultimately results in ROS production and DNA damage.^26^ Therefore, our aim was to investigate whether the combined treatment induces cellular death through an excessive ROS production. The cells underwent treatment with NCX 4040 at 5 μM and Napabucasin at 100 and 250 nM, both individually and in combination. Flow cytometry was utilized to analyze both cellular and mitochondrial ROS levels. As depicted in Figure 4 the combination of NCX 4040 and Napabucasin led to a significant increase in both cellular (see Figure 4A-4D) and mitochondrial ROS levels (see Figure 4E–4H), surpassing their individual effects. This implies that one plausible mode of cell death is through oxidative stress, necessitating further investigation.

**Figure 4:**
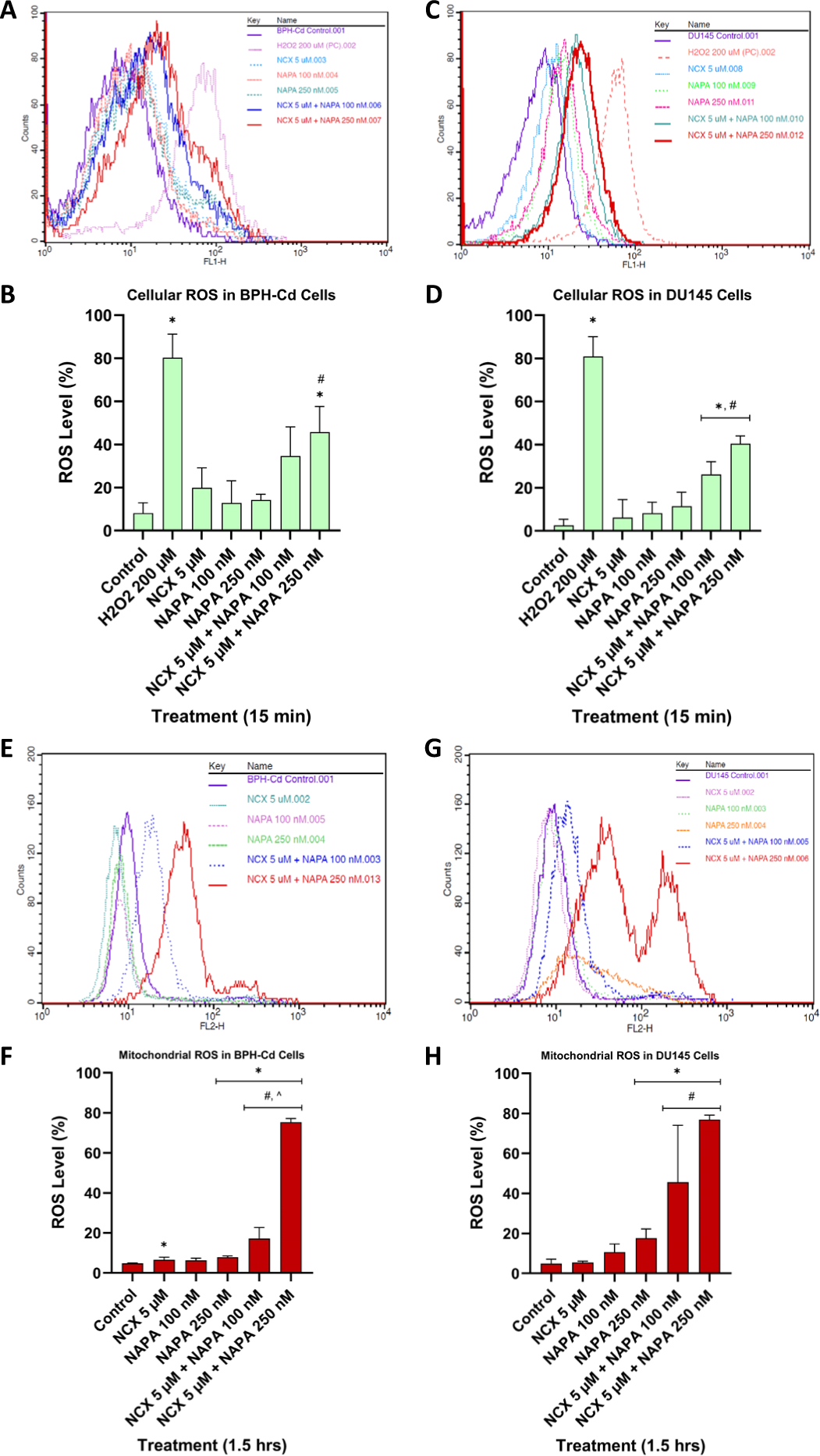
The combination of NCX 4040 and Napabucasin induces robust cellular (top panel) and mitochondrial (bottom panel) ROS production. The DCFH-DA ROS assay was employed to determine the extent of cellular ROS production upon combination treatment. Upon 60-70% confluency, the cells were treated with 20 μM of DCFH-DA followed by respective drug treatments. The mitoSOX red ROS assay was performed to detect mitochondrial ROS production upon combination treatment. Remarkably, the combination treatment elicited a substantial rise in the percentage of both cellular and mitochondrial ROS compared to the individual treatments, suggesting that the treatment induces oxidative stress-related cellular death. A positive control of hydrogen peroxide was used to validate the assay. **A)** and **C)** refers to the flow cytometric histograms for cellular ROS and **E)** and **G)** refers to histograms for mitochondrial ROS. **B), D), F)** and **H)** represent the subsequent quantitative analysis. Data are means ± SD (N=3). * p < 0.05: compared to control; and # p < 0.05: mean of combination treatment compared to their respective individual treatments.

Expanding our inquiry, we explored the potential of this combined treatment in enforcing cell cycle arrest. Our findings demonstrated that the combination of NCX 4040 and Napabucasin instigated G2/M cell cycle arrest in both cell lines by 95% in BPH-Cd and 70% in DU145 cells, preventing mitosis and subsequent cell proliferation. In stark contrast, individual treatments failed to elicit any such discernible impact, as shown in Figure 5A and 5B.

**Figure 5:**
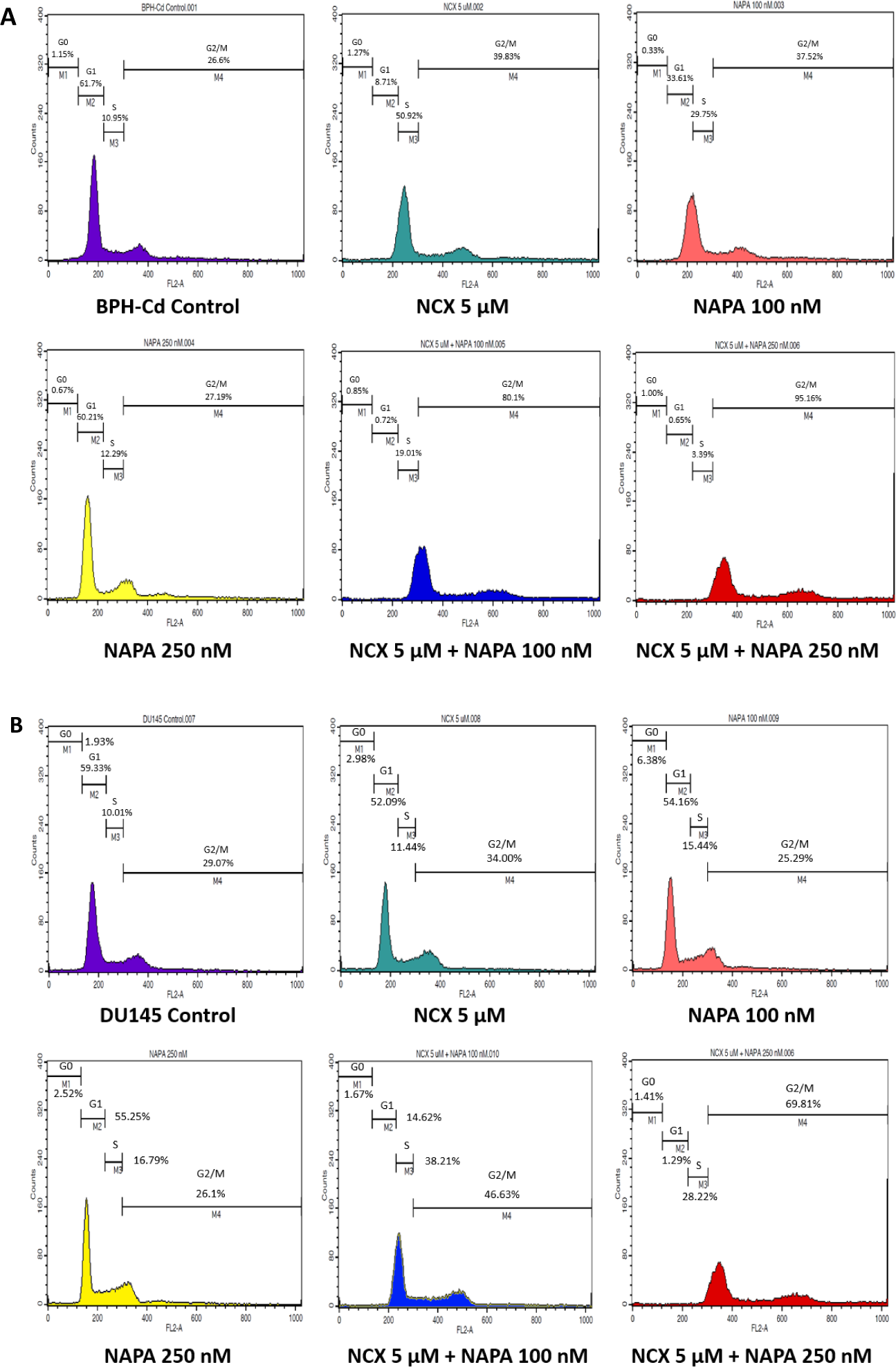
NCX 4040 and Napabucasin, in combination, induces G2/M cell cycle arrest in A) BPH-Cd and B) DU145 cells. Cell cycle analysis was performed to analyze whether combination treatment could enforce cell cycle arrest. Upon 60-70% confluency, the cells were subjected to varying concentrations of NCX 4040 and Napabucasin, both individually and in combination, for a duration of 48 hours. The flow cytometric technique was employed to scrutinize the DNA content of the cells, and we observed a notable shift towards the G2/M peak in the combination treatment, indicative of G2/M cell cycle arrest. Our findings suggest that the combination treatment impedes subsequent mitosis and cell proliferation, as illustrated by the considerable G2/M peak shift. Data is representative of one of three experiments (N=3). Significant differences were observed between control and treatment groups and individual and combination treatment groups determined by Dunnett’s and Tuckey’s statistical test, respectively.

To ascertain the type of cell death, we employed the FITC Annexin V/PI apoptosis detection kit, facilitating the detection of necrotic, early, and late apoptotic cell death. As illustrated in Figure 6A and 6B, the combination treatment induced late apoptosis in both BPH-Cd and DU145 cells, unlike individual treatments, which had no significant effects. Our results indicate that the combined treatment holds the potential to trigger apoptosis in cancer cells, thereby eliminating them.

**Figure 6:**
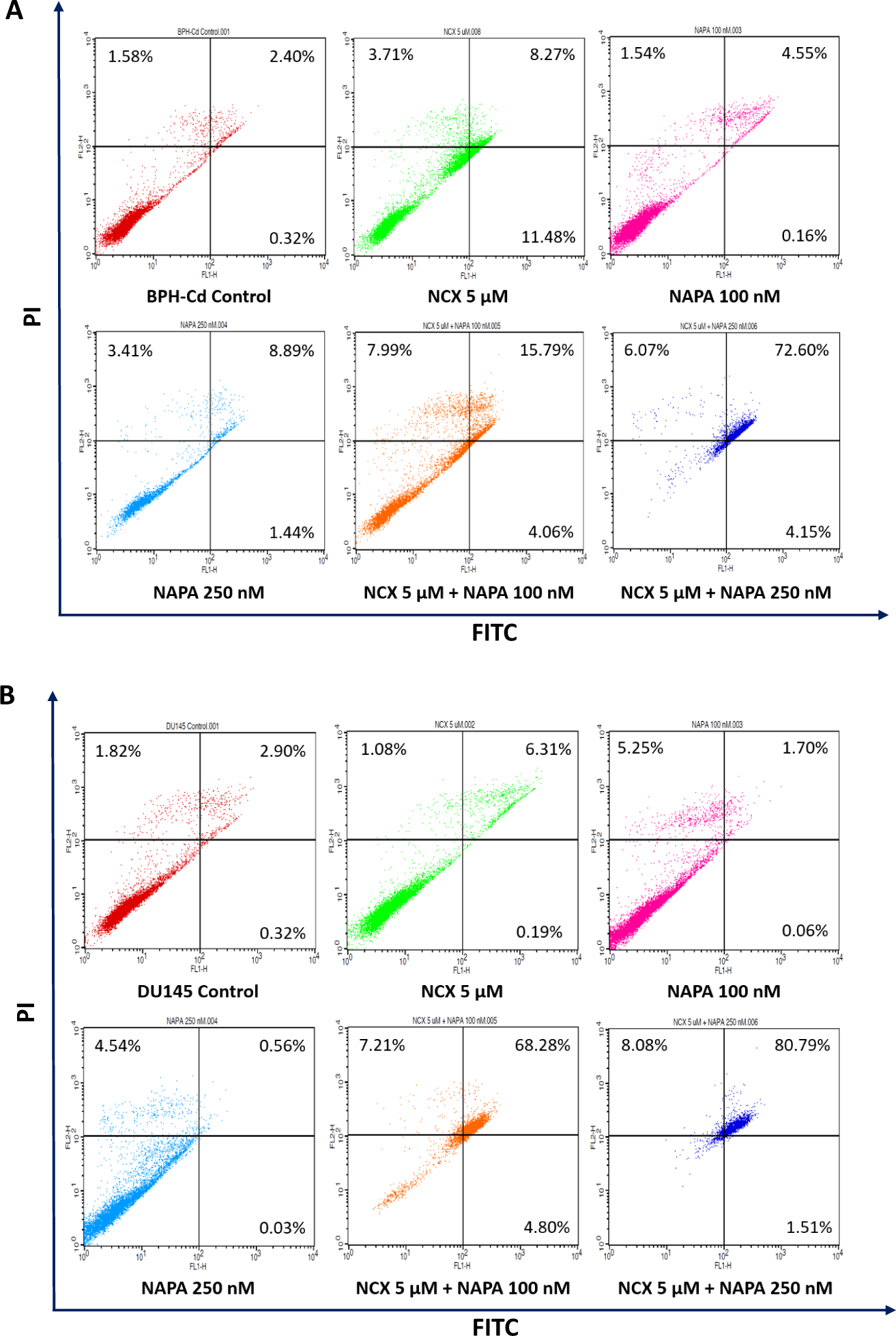
The combination of NCX 4040 and Napabucasin causes late apoptosis in A) BPH-Cd and B) DU145 cells. Cell death analysis was performed to determine the type of cell death. When the cells were 60% confluent, they were serum starved overnight, followed by treatment for 48 hours. Later, the cells were collected and stained using FITC Annexin V Apoptosis kit. Data depicts that the combination treatment induces significant apoptosis as compared to individual treatments. Data is representative of one of three experiments (N=3).

### Combination treatment regulates the expression of key markers in STAT3 and redox homeostasis and signaling pathway

Based on our flow cytometric analysis and cell viability assay, it is unambiguous that the combination treatment exerts a synergistic effect on inducing cellular and mitochondrial ROS production in both BPH-Cd and DU145 cells. Moreover, a comprehensive review of the literature provides valuable insights into the pivotal role of STAT3 downregulation or inhibition. Consequently, we conducted a western blot analysis to examine several markers from the STAT3 and redox homeostasis and signaling pathway.

As shown in Figures 7A and 7B, we observed a remarkable and distinct downregulation of both cytosolic (pSTAT3–705) and mitochondrial (pSTAT3–727) STAT3 in response to the combination treatment in both cell lines, in stark contrast to the effects of individual treatments. As highlighted in numerous studies, STAT3 is critically implicated in the regulation of carcinogenesis, chemoresistance, and modulation of cancer stemness. Persistent upregulation of STAT3 fosters uncontrolled cell growth and progression of the disease.^35^ Therefore, our study provides compelling evidence that the combination treatment triggers STAT3 downregulation impeding proliferation, growth, and migration.

**Figure 7:**
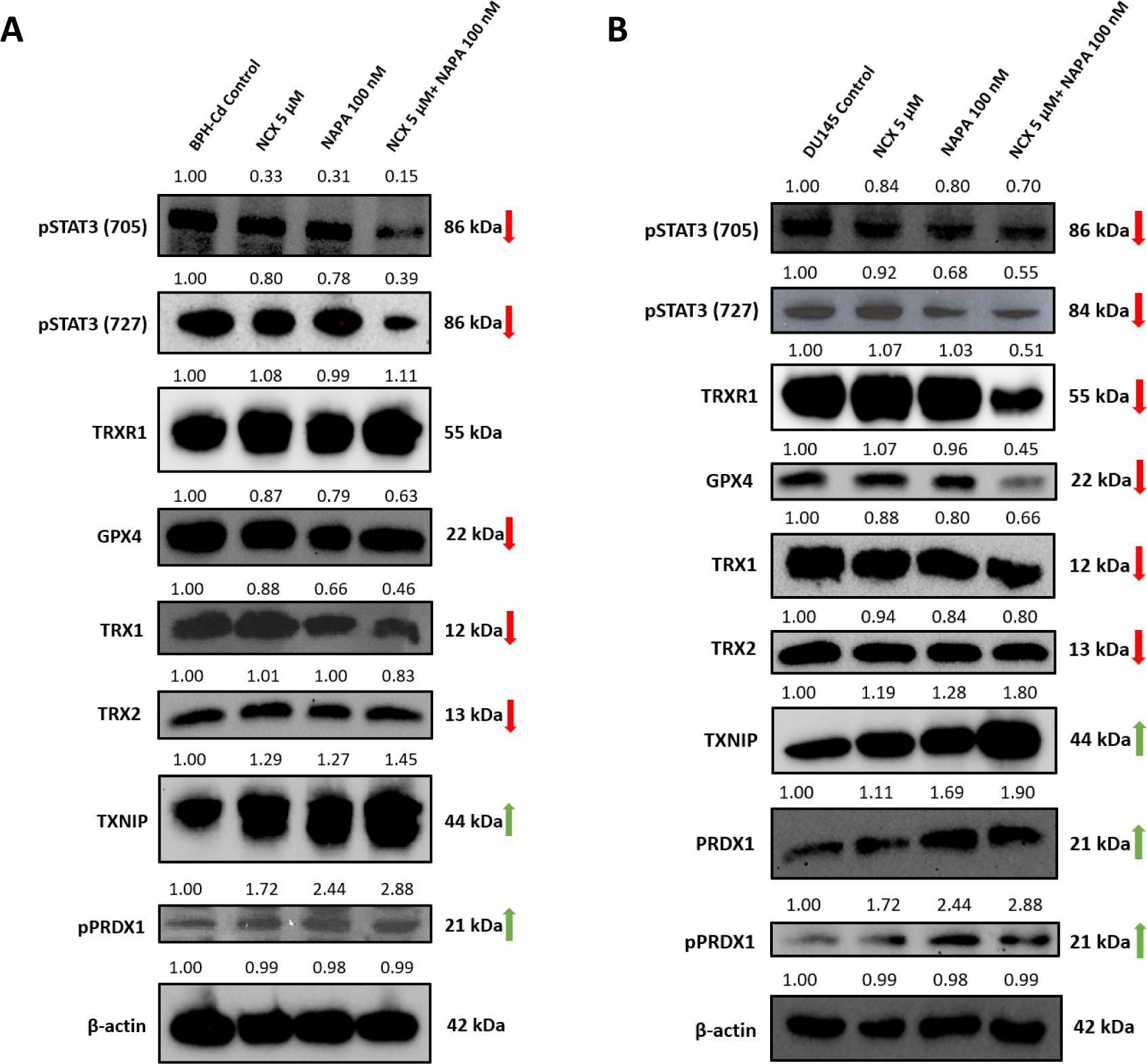
Combination treatment regulates the expression of key markers in STAT3 and redox homeostasis and signaling pathway. A Western blot analysis was performed to evaluate the expression of key markers involved in STAT3 and redox homeostasis and signaling pathway **(A and B)**. Cytosolic (pTyr 705) and mitochondrial (pSer 727) STAT3, TRXR1, GPX4, TRX1 and 2 were highly downregulated. On the other hand, TXNIP and pPRDX1 were highly upregulated in both the cell lines. This indicates that the combination treatment is inhibiting cell proliferation and growth and causing oxidative stress related cellular death.

Furthermore, evaluating the expression of key targets from the redox homeostasis and signaling pathway resulted in the downregulation of thioredoxin (TRX1/2), thioredoxin reductase 1 (TRXR1), and glutathione peroxidases (GPX4)—crucial regulators maintaining ROS homeostasis and detoxification.^36^ Moreover, cells treated with the combination drug exhibited a significant upregulation of thioredoxin-interacting protein (TXNIP) and peroxiredoxins (PRDX1) within the pathway, elucidating the elevated ROS production in both cell lines and corroborating the results from the flow cytometric analysis shown in Figure 4.

In summary, our study underscores the remarkable potential of the combined treatment in inducing ROS production and modulating the STAT3 signaling pathway.

### NCX 4040 and Napabucasin in combination decreases the expression of key cancer stem cell-like markers

Cancer comprises a heterogeneous population of cells, including a distinct subset exhibiting stem cell-like characteristics, known as cancer stem cells (CSCs). The eradication of CSCs is crucial for effective cancer therapy, as these cells can differentiate into cancer cells, leading to tumor relapse, and drive epithelial-to-mesenchymal transition (EMT), contributing to metastasis.^37^ As previously discussed, Napabucasin (BBI608) has been reported as an effective inhibitor of cancer stemness by targeting downstream signaling of the JAK-STAT pathway. In alignment with this, our findings demonstrate that the combination therapy of NCX 4040 and Napabucasin downregulates STAT3 signaling markers, which are known regulators of cancer stemness. To further investigate the effects of this combination therapy on cancer stemness, we evaluated its impact on two key CSC-associated signaling pathways: the Delta-Notch and Wnt/β-catenin pathways.

As shown in Figures 8C and 8D, the combination therapy significantly reduced the expression of key markers involved in the Delta-Notch and Wnt/β-catenin signaling pathways, both of which play critical roles in CSC maintenance and tumor progression. Notch signaling is essential for cancer cell survival, while the crosstalk between the Wnt/β-catenin pathway and the androgen receptor (AR) enhances AR transcriptional activity, even at low androgen levels, contributing to the progression of castration-resistant prostate cancer (CRPC).^38–39^ Additionally, we observed a marked decrease in the transcription factors SOX-2, OCT-4, and Nanog, which are crucial for maintaining cancer stemness and function as downstream effectors of these pathways.

**Figure 8:**
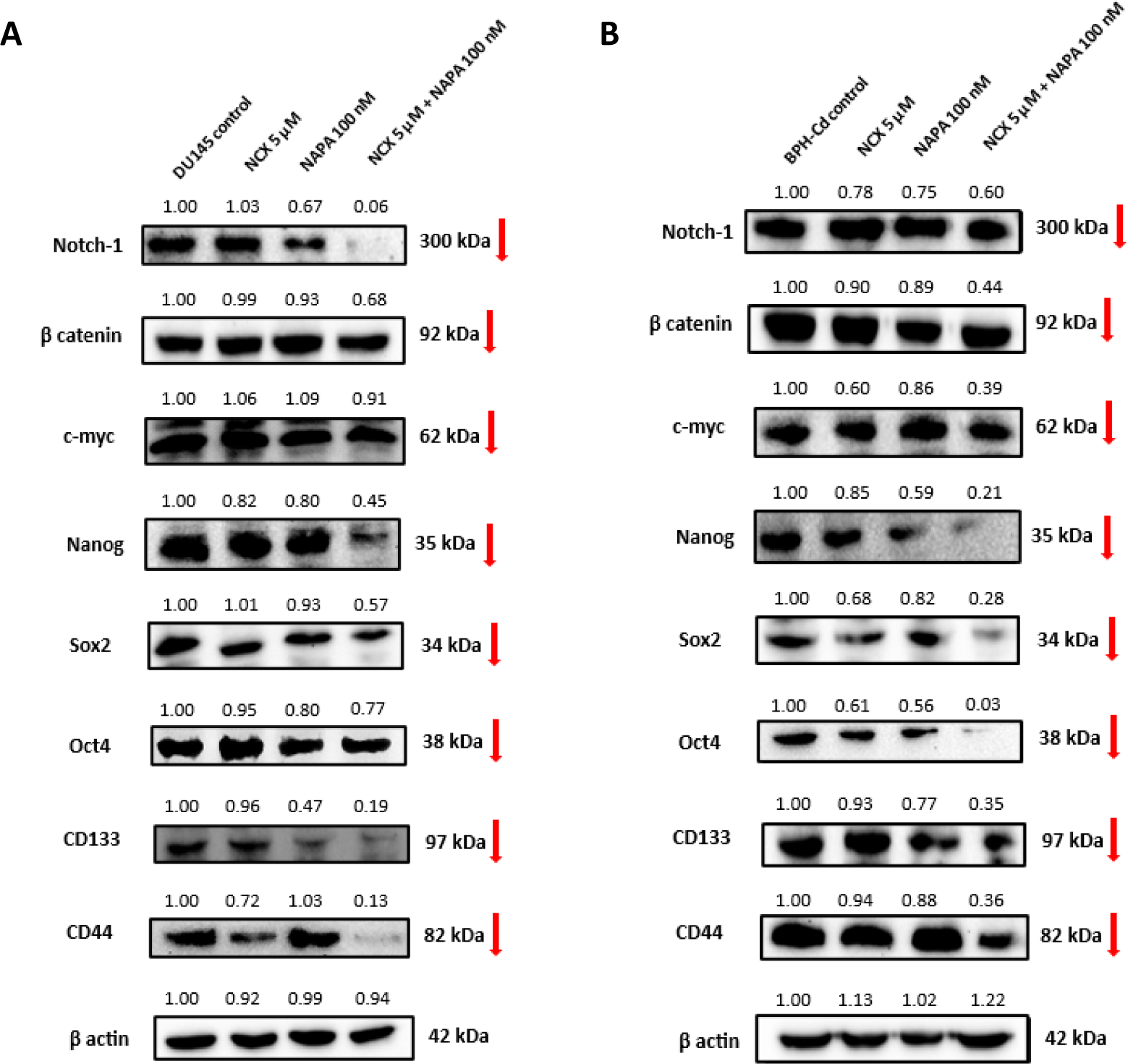
NCX 4040 synergizes with Napabucasin to downregulate cancer stemness markers in both A) BPH-Cd and B) DU145 cells. Western blot analysis of these cells demonstrates a significant reduction in Notch-1 and β-catenin—key regulators of distinct stemness signaling pathways—along with the downregulation of transcription factors such as Sox2, Oct4, Nanog, and c-Myc, as well as stem cell surface markers CD133 and CD44. These findings highlight the synergistic effect of the combination treatment in diminishing the stemness characteristics of PCa cells.

Furthermore, the combination treatment led to a significant reduction in the expression of CSC-associated surface markers CD44 and CD133, which are predominantly found in cancer stem cells. This effect was consistently observed across both PCa cell lines tested. Collectively, these findings suggest that the combination therapy effectively weakens CSC characteristics in aggressive PCa models, underscoring its potential as a promising therapeutic strategy for targeting CSC-driven tumor progression.

### The combination treatment increases the expression of DNA damage marker, pH2AX

To evaluate whether the combination treatment of NCX 4040 and Napabucasin induces DNA damage, we assessed the expression of phosphorylated histone H2AX (pH2AX), a well-established marker of DNA double-strand breaks (DSBs).^40^ Given that excessive reactive oxygen species (ROS) can contribute to genomic instability and activate apoptotic pathways, we hypothesized that the increased ROS levels observed with the combination treatment would correlate with elevated DNA damage.

Immunofluorescence analysis revealed a significant increase in nuclear pH2AX foci in BPH-Cd cells following combination treatment (Figure 9). This increase was markedly higher in cells treated with NCX 4040 (5 μM) in combination with Napabucasin (100 nM or 250 nM) compared to individual treatments, suggesting a synergistic effect in inducing DNA damage. Notably, cytoplasmic spillover and accumulation of pH2AX foci was observed, indicating persistent DNA damage likely resulting from oxidative stress exceeding cellular DNA repair capacity.

**Figure 9:**
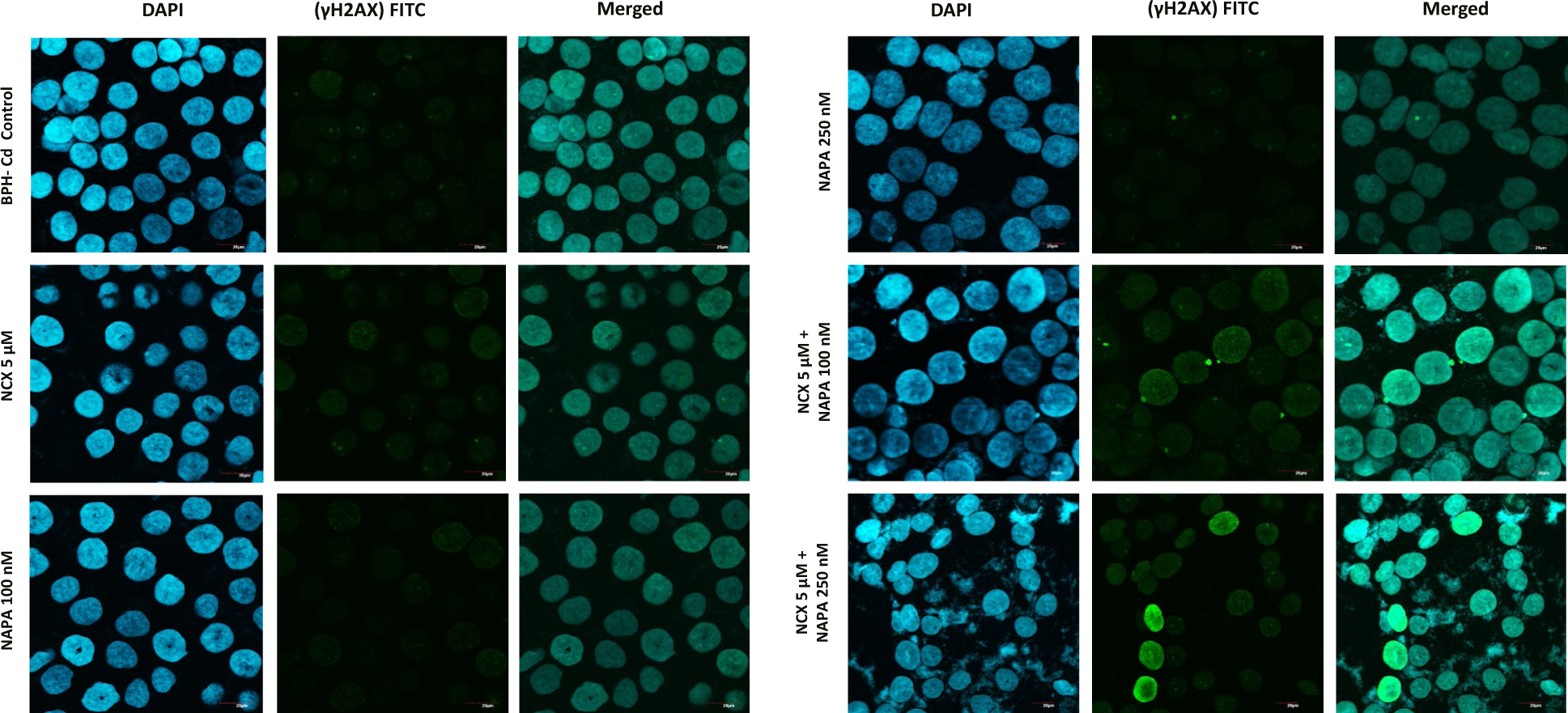
Combination therapy with NCX 4040 and Napabucasin induces elevated pH2AX expression in BPH-Cd cells. Immunofluorescence analysis of BPH-Cd cells treated with NCX 4040, Napabucasin, and their combination reveals the highest pH2AX expression in the combination treatment group, as indicated by significantly greater fluorescence intensity in the FITC channel compared to both the control and individual treatment groups. DAPI staining was used to visualize DNA content, and merged images of the DAPI and FITC channels were analyzed to localize pH2AX expression. Images were captured at 60× magnification.

These findings are consistent with our previous results demonstrating late-stage apoptosis induction and ROS production (Figures 4 and 6), further supporting the notion that combination therapy promotes cell death via excessive ROS generation and subsequent genomic instability. Collectively, our data provides strong evidence that NCX 4040 and Napabucasin act synergistically to enhance DNA damage, highlighting their potential as an effective therapeutic strategy for PCa.

### NCX 4040 and Napabucasin regulates the expression of key markers in cGAS-STING antitumor immunogenic pathway

The cyclic guanosine monophosphate-adenosine monophosphate synthase (cGAS) stimulating interferon gene (STING) antitumor immunogenic pathway is an evolutionary conserved cytosolic DNA sensor pathway that can initiate innate immune defense responses produced by type I interferon and other immune mediators.^41^ Our findings from immunofluorescence imaging, aimed at assessing DNA damage, unveiled the emergence of nuclear release and increased expression of pH2AX 48 hours after treatment. These noteworthy observations prompted our inquisition to perform a screening study on cGAS-STING pathway.

As illustrated in Figures 10, all downstream markers of the pathway, including cGAS, STING, tank binding kinase 1 (TBK1), and interferon response factor 3 (IRF3), exhibited remarkable upregulation in response to the combination treatment, in contrast to the individual treatment groups. These intriguing results strongly suggest that NCX 4040 and Napabucasin possess the capability to stimulate the antitumor immunogenic pathway, potentially leading to T cell priming and infiltration. However, it is important to note that these results are limited to screening these proteins, and a more in-depth investigation is currently underway to assess their immunotherapeutic potency in-vivo.

**Figure 10:**
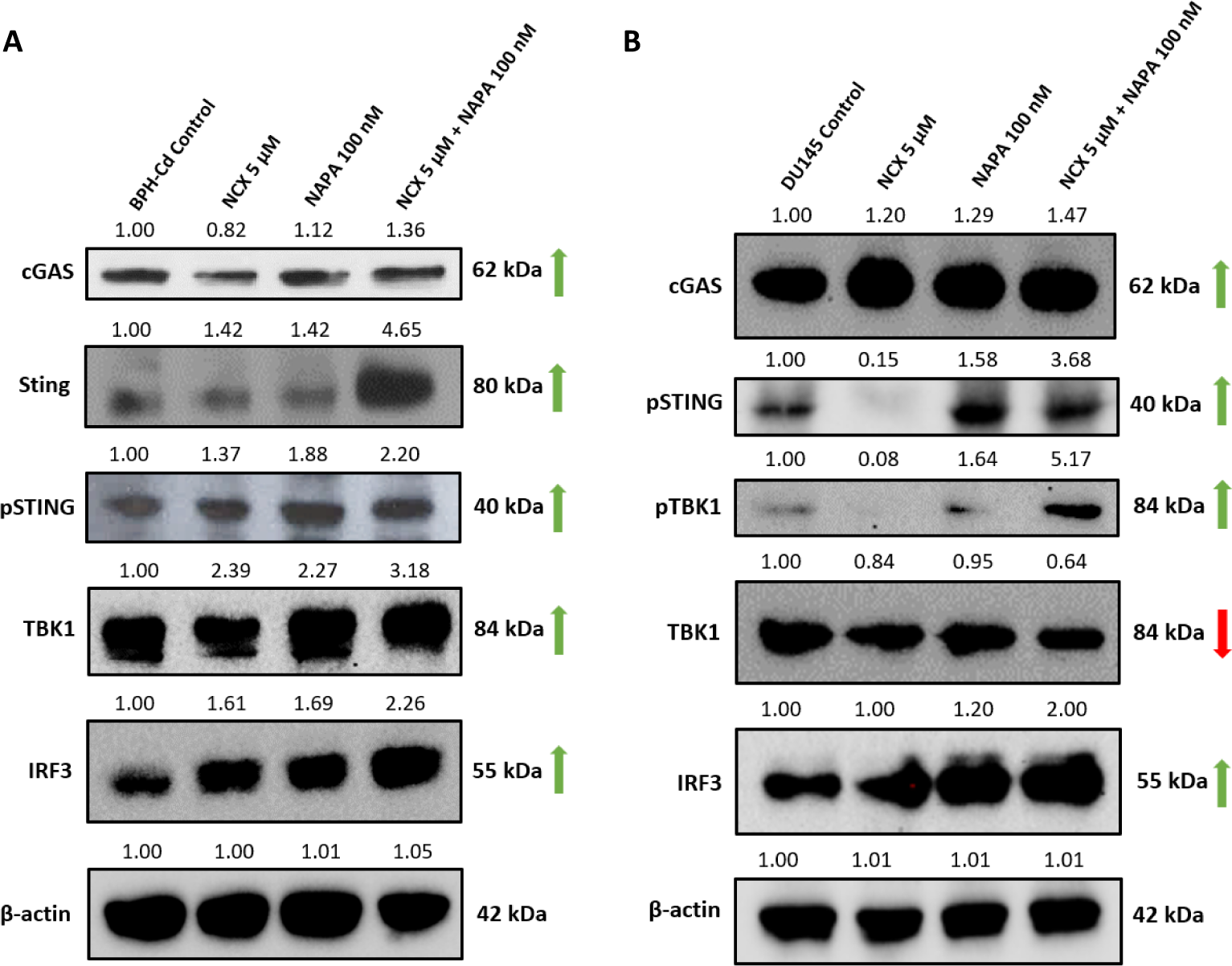
The combination treatment activates antitumor immunogenic cGAS STING signaling both the cell lines (A and B). All the downstream markers in the pathway are highly upregulated, indicating at antitumor immunogenic response. This implies that the combination treatment might lead to type I interferon response and T cell priming, resulting in calling of the immune system.

## Discussion

PCa is the most common non-cutaneous malignancy in American men and the second leading cause of cancer-related morbidity after lung cancer.^42^ Annually, about 1.3 million cases are diagnosed worldwide, with approximately 1 in 8 men being diagnosed each year.^1^ Current treatments for advanced PCa - chemotherapy, hormone therapy, radiation, and surgery - are often inadequate due to severe side effects, toxicity, chemoresistance, cancer stemness and limited improvement in survival rates. These treatments are effective for localized diseases but often fail in high-risk or metastatic cases. The heterogeneity and diverse etiology of PCa necessitates a multifaceted therapeutic approach.

To target some of the complex attributes of PCa discussed in this study namely STAT3, generation of cellular and mitochondrial oxidative stress, DNA damage and antitumor immunogenic response, monotherapeutic approaches remain less potent and obsolete. The combinatorial therapy approach has emerged as a useful treatment option for various types of cancer to overcome the shortcomings of existing treatment modalities. Therefore, in this study we explored the synergy and combinatorial effects between two drug compounds, namely NCX 4040 and Napabucasin. While individual efficacy of these drugs has been shown in other cancers, their combined effect on advanced PCa is largely not explored.

Combining NCX 4040 with Napabucasin demonstrated synergistic anti-cancer effects in BPH-Cd and DU145 cells, reducing cell viability, tumorigenic potential, and metastatic properties more effectively than individual treatments. The combination treatment showed dose-dependent inhibition of cell viability and enhanced anti-cancer effects, supported by synergy scores indicating effective hindrance of cell proliferation and growth. The less toxic effect on benign prostate cells suggested reduced off-target cytotoxicity.

Further assays confirmed the combination’s superior suppression of tumorigenic potential, evidenced by reduced colony formation, spheroid size, and migratory capabilities, suggesting significant inhibition of in-vitro tumor formation and potential in mitigating metastatic spread in advanced PCa management. However, structural and pharmacodynamic studies are needed to fully understand the synergy mechanism, and further strategies are required to enhance targeting in clinical settings.

Further experimentation revealed the mechanism of action for the combination therapy. Other studies have underscored the potential of NCX 4040 to induce oxidative stress by depleting cellular glutathione (GSH), resulting in the generation of reactive oxygen species (ROS) or quinone methide, culminating in DNA damage and cell death.^10^ On the other hand, Napabucasin have been shown to target STAT3 signaling, self-renewal in cancer and immunogenic cell death in many cancer cell types.^15,26,29^ Hence, we were interested to see if the combination could trigger such responses and target multiple aspects in PCa leading to cellular demise. Flow cytometric analysis showed high levels of cellular and mitochondrial ROS, especially in combination-treated cells, indicating oxidative stress as the primary mode of cell death, supported by western blot results.

STAT3 is crucial in tumorigenesis, influencing gene transcription related to survival, apoptosis resistance, metastasis, and angiogenesis.^43^ STAT3 is also redox-sensitive, with cysteine residues susceptible to oxidation. Post-translational modifications (PTMs) induced by ROS are known to impact on the activity or stability of STAT3.^44^ Our results demonstrated a significant reduction in both cytosolic Tyr705 and mitochondrial Ser727 phosphorylated isoforms of STAT3 upon exposure to the combination of NCX 4040 and Napabucasin, highlighting the impedance of cancer progression. TRX1/2, TRXR1, and GPX1/4 are integral pro-survival proteins within the redox pathway, which also facilitate the phosphorylation of STAT3 at Tyr705 to enable dimerization and its transcriptional activity. Studies have demonstrated that inhibition of these proteins results in the accumulation of oxidized PRDX1 and TXNIP, thereby elevating oxidative stress and blocking STAT3-mediated transcription, ultimately impeding cell proliferation and survival.^45^ Hence, the downregulating effect of the combination treatment on these potent enzymes proves to be highly effective in inducing oxidative stress.

It is important to note that TXNIP plays a dual role. Not only does it mediate oxidative stress, but it also functions as a potential tumor suppressor gene, effectively inhibiting the proliferation, invasion, and migration of tumor cells while promoting apoptosis.^46^ Therefore, the noteworthy upregulation of TXNIP observed in our study represents a novel and promising advancement in the pursuit of targeting PCa. In addition to TXNIP, PRDX1 also plays an important role in tumor suppression, apoptosis, and inhibition of cell growth and proliferation. PRDX1 has recently emerged as H_2_O_2_ scavenger, however, in presence of excessive H_2_O_2_, PRDX1 can self-oxidize leading to oxidative stress.^47^ This compellingly suggests that the synergistic interplay between NCX 4040 and Napabucasin has a pronounced effect on crucial cancer survival and progression participants.

Furthermore, cell cycle analysis suggested that the combination of NCX 4040 and Napabucasin causes G2/M cell cycle arrest, effectively disrupting subsequent mitosis and thereby limiting cell proliferation. Cell death analysis corroborated this finding by demonstrating a high percentage of late apoptotic cells, indicative of cell death occurring when cells reach the end of their natural lifespan. In contrast, no significant cell cycle arrest or cell death was observed in cells treated with either drug individually.

Drawing upon existing literature, we recognized Napabucasin’s potential to induce DNA damage and NCX 4040’s ability to trigger oxidative stress-induced cell death. Building on this background and our previous results, we performed immunofluorescence imaging with pH2AX to investigate the effects of the combination treatment on DNA damage. H2A.X is essential for checkpoint-mediated cell cycle arrest and DNA repair following double-stranded DNA breaks. During apoptosis, DNA damage and fragmentation lead to the rapid phosphorylation of H2A.X, which inhibits the recruitment of DNA repair proteins and promotes the binding of pro-apoptotic factors, ultimately causing cell death.^40^ Observing a high percentage of late apoptotic cells and G2M cell cycle arrest, we hypothesized an increased expression of H2A.X in the combination treatment, as confirmed in Figure 7.

While the combination treatment for PCa shows great promise in producing chemotherapeutic effects, the complex nature of cancer demands a multifaceted approach. It is crucial to develop strategies that harness the body’s antitumor immunogenic response to combat the disease effectively. Activating this response can help preempt the evasive maneuvers of malignant cells and reduce disease progression, ensuring optimal treatment outcomes for patients.

Our immunofluorescence imaging data revealed novel insights into the dynamic interplay between NCX 4040 and Napabucasin and prompted us to investigate the cGAS-STING pathway. This pathway triggers an immune response against tumor cells by detecting and responding to DNA damage, leading to cytokine production and immune cell recruitment to the damage site.^41^ Western blot analysis showed that the combination of NCX 4040 and Napabucasin upregulated key players in the cGAS-STING pathway. The downstream molecule, IRF3, was notably upregulated in both BPH-Cd and DU145 cell lines, suggesting that the combination treatment could trigger an antitumor immune response, a crucial component of successful cancer therapy. This finding indicates that our combination treatment not only generates chemotherapeutic effects via oxidative stress but also stimulates an immunotherapeutic effect, potentially activating the immune system to target advanced PCa. To be clinically relevant, further experimentation in relevant mouse models is imperative, and our preliminary investigation is measuring T cell activity.

Our data strongly support the enhanced anticancer efficacy of the drug combination in inhibiting key drivers of PCa. In addition to these effects, we investigated the impact of combination therapy on cancer stemness, a critical contributor to tumor progression, metastasis, recurrence, and therapy resistance. Our findings demonstrated a significant downregulation of two essential CSC signaling pathways, Notch-1 and Wnt/β-catenin, in response to treatment.

Notch-1 signaling is crucial for prostate gland development during organogenesis but remains largely quiescent in adult tissues. Aberrant activation of this pathway promotes luminal cell proliferation, contributing to tumorigenesis. Notch-1 is significantly expressed in PCa cell lines, including LNCaP, DU145, PC3, and C4-2B, where it regulates the expression of stemness-associated genes such as Sox2, c-Myc, Oct4, Nanog, and CD133^48–50^ Sox2 and Oct4 are pivotal for CSC self-renewal and proliferation, while Nanog regulates differentiation.^51–52^ Our results indicate that combination therapy effectively downregulates Notch-1 signaling, leading to reduced expression of these key markers and, consequently, suppression of cancer stemness in PCa cells.

Similarly, Wnt/β-catenin signaling is well established as a driver of PCa progression, tumor development, and castration resistance. Activation of this pathway leads to the upregulation of oncogenic targets, particularly c-Myc, which plays a central role in tumor growth and metabolic reprogramming. Our findings indicate that combination therapy significantly downregulated both β-catenin and c-Myc, as well as the cell cycle regulator Cyclin D, suggesting a broader inhibitory effect on tumor proliferation.^38–39^ While Wnt signaling has been extensively studied across various malignancies, its precise role and downstream mechanisms in PCa remain incompletely understood. Our study provides further evidence supporting the therapeutic potential of targeting this pathway.

Beyond individual pathway inhibition, we also examined the impact of combination therapy on CSC populations by evaluating the expression of CD44 and CD133, two well-recognized cell surface CSC markers. The observed downregulation of these markers suggests a significant reduction in CSC populations, indicating a potential to mitigate tumor recurrence and therapeutic resistance.

In conclusion, our innovative combination regimen marks a significant advancement in cancer treatment, eliciting a robust chemoimmunotherapeutic response against advanced PCa. By effectively targeting cancer stemness, this approach holds promise not only in enhancing therapeutic efficacy but also in weakening the cancer stem cell compartment—an essential factor for achieving sustained disease control in advanced PCa.

### Proposed mechanism of action

Based on our findings in this study, we have successfully demonstrated that the administration of NCX 4040 in combination with Napabucasin effectively thwarts the insidious expression of STAT3 and subsequently induces a cascade of oxidative stress-mediated cellular apoptosis and DNA damage. This remarkable phenomenon culminates in the momentous activation of cGAS STING pathway, which remarkably provokes an anti-tumor immunogenic response.

Notably, our intricate mechanistic elucidation revealed that the combination treatment’s powerful efficacy is attributed to its robust suppression of both cytosolic and mitochondrial STAT3 by proficiently targeting the pro-survival redox markers TRX1/2, TRXR1, and GPX4 while simultaneously upregulating redox promoters TXNIP and PRDX1. These salient attributes unequivocally underscore the extraordinary chemotherapeutic potential of this synergistic amalgamation.

Moreover, our research further reveals that this combination therapy also serves as a potent immunotherapeutic agent by inducing an anti-tumor immunogenic response via the activation of cGAS-STING pathway. This engenders cancer cells to be exposed to the immunosurveillance system and ultimately culminates in the priming of T cells and galvanizing of the immune system.

To explore subtly towards the cancer stemness of PCa, significant downregulation in STAT3 signaling pathway speculated the downregulation of major stemness related pathways. Two proteins Notch-1 and β catenin have shown downregulation. In addition, downregulation of CSC related transcription factors and cell surface markers adds on to the provided evidence. This indicates that two major stemness related pathways Delta-Notch signaling and Wnt-β catenin signaling pathways are hindered.

A concise proposed mechanism of action is depicted in Figure 11.

**Figure 11:**
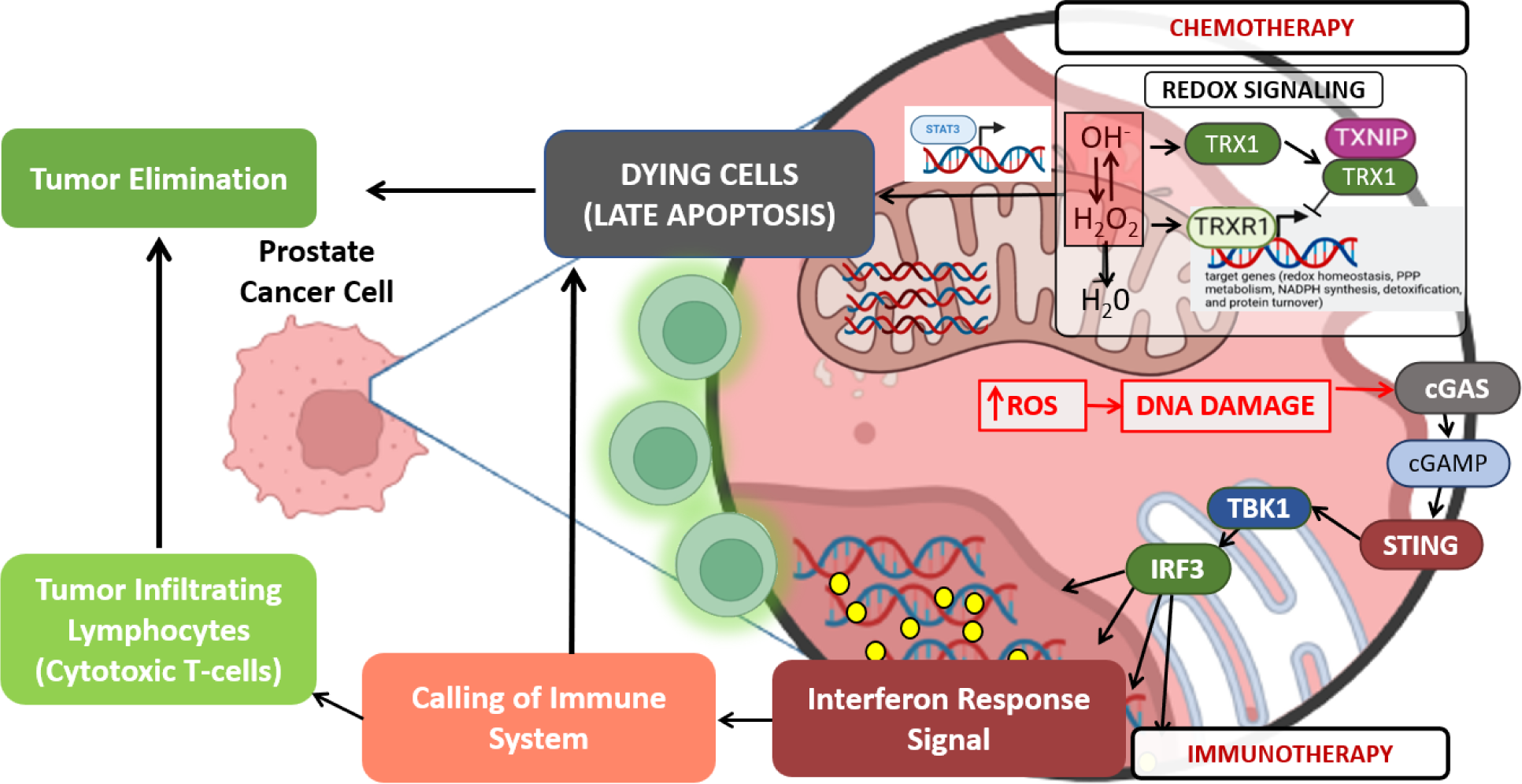
Proposed mechanism of action of the synergistic combination of NCX 4040 and Napabucasin in targeting advanced PCa. Synergistic drug therapy triggers chemoimmunotherapeutic response by inducing oxidative stress via disrupting the redox homeostasis signaling and STAT3 inhibition, inducing DNA damage, and thereby activating the anti-tumor immunogenic pathway, cGAS-STING.

## Conclusion

Our groundbreaking study presents a potent solution for advanced PCa treatment by combining NCX 4040 and Napabucasin. This chemoimmunotherapeutic approach significantly reduces cell growth, proliferation, tumorigenicity, cancer stemness and metastasis in cadmium-transformed and advanced PCa cells. The therapy induces cellular and mitochondrial ROS production, leading to late apoptosis and G2/M cell cycle arrest, and disrupts redox homeostasis by altering key protein levels involved in detoxification pathways. Additionally, the treatment triggers immunogenic activation of the cGAS-STING pathway, oxidative stress, and STAT3 inhibition, potentially overcoming cancer cell immune evasion. Despite some limitations, our findings suggest this combination therapy holds great promise as a novel PCa treatment, offering hope for improved patient outcomes and further research opportunities.

## Acknowledgments

The authors would like to thank Dr. Weihai Zhan, Director of Student Research, Office of Research, University of Illinois College of Medicine, for conducting a comprehensive statistical analysis of our data.

Additional Information:

## Author’s Information

**Full name:** Nishtha Pathak (N.P.)
**Mailing address:** 30 Gardner Rd, Apt 2E, Brookline, MA - 02445

**Full name:** Shrushti Shah (S.S.)
**Mailing address:** 914 West Blvd, Hartford, CT 06105

**Full name:** Jeffrey Mathew (J.M.)
**Mailing address:** 1552 Oakforest Drive, Rockford, IL, 61107

**Full name:** Gnanasekar Munirathinam (G.M.)
**Mailing address:** 1601 Parkview Ave, Rockford, IL 61108

## Conflicts of Interest

The authors declare no potential conflicts of interest. The funders had no role in the design of the study, in the collection, analyses, or interpretation of data, in the writing of the manuscript, or in the decision to publish the results.

## Data Availability Statement

The data represented in the paper are available on request from the corresponding author.

## Ethics Statement

NA

## Funding

The study is partly supported by Master’s in Medical Biotechnology program, Department of Biomedical Sciences, University of Illinois College of Medicine, Rockford, IL, USA – 61108.

## Author Contributions

Conceptualization: N.P. and G.M.; methodology: N.P., S.S., and J.M.; formal analysis: N.P., and S.S.; investigation: N.P., and S.S.; resources: G.M.; writing—original draft preparation: N.P., S.S. and G.M.; writing—review: N.P. and G.M.; supervision: G.M.; project administration: G.M.; funding acquisition: G.M. All authors have read and agreed to the published version of the manuscript.

## Notes

### Competing Interest Statement

The authors have declared no competing interest.

